# DDX21 is a p38-MAPK sensitive nucleolar protein necessary for mouse preimplantation embryo development and cell-fate specification

**DOI:** 10.1101/2021.04.13.439318

**Authors:** Pablo Bora, Lenka Gahurova, Andrea Hauserova, Martina Stiborova, Rebecca Collier, David Potěšil, Zbyněk Zdráhal, Alexander W. Bruce

## Abstract

Successful navigation of the mouse preimplantation stages of development, during which three distinct blastocyst lineages are derived, represents a prerequisite for continued development. We previously identified a role for p38-mitogen-activated kinases (p38-MAPK) regulating blastocyst inner cell mass (ICM) cell-fate, specifically primitive endoderm (PrE) differentiation, that is intimately linked to rRNA precursor processing, polysome formation and protein translation regulation. Here, we develop this work by assaying the role of DEAD-box RNA helicase 21 (*Ddx21*), a known regulator of rRNA processing, in the context of p38-MAPK regulation of preimplantation mouse embryo development. We show nuclear *Ddx21* protein is robustly expressed from the 16-cell stage, becoming exclusively nucleolar during blastocyst maturation; a localisation dependent on active p38-MAPK. Efficient siRNA mediated clonal *Ddx21* knockdown within developing embryos is associated with profound cell autonomous and non-autonomous proliferation defects and reduced blastocyst volume, by the equivalent peri-implantation blastocyst stage. Moreover, ICM residing *Ddx21* knockdown clones express the EPI marker NANOG but rarely express the PrE differentiation marker GATA4. These data contribute extra significance to emerging importance of lineage specific translation regulation, as identified for p38-MAPK, during mouse preimplantation development.

## Introduction

Mammalian preimplantation embryonic development is the period between fertilisation and uterine implantation. This period encompasses zygotic genome activation (ZGA), cellular proliferation and lineage specification, culminating in a metabolically active outer trophectoderm (TE) cell layer, fluid-filled blastocoel cavity, epithelial primitive endoderm (PrE) layer and pluripotent epiblast cells (EPI) enveloped between the TE and PrE. Primary transcriptional regulators of cell-fate, such as TEAD4 and CDX2 (TE fate), GATA6, SOX17 and GATA4 (PrE fate) and NANOG and SOX2 (EPI fate), have been identified in the mouse model and are enabled by specific signalling networks; *i*.*e*., HIPPO-YAP for TE and FGF4-FGFR1/2-MEK/ERK for PrE (as extensively reviewed recently^1–4^). Furthermore, mechanical forces acting on the developing early embryo, on both cellular and embryonal levels, have been found to be significant in the spatial sorting and cell-fate specification of individual blastomeres^5–7^. Additionally, a more holistic view of preimplantation embryonic development is emerging by the incorporation of regulatory studies targeting global events, such as translation^8–11^. Indeed, regulation of translation is reported to be vital to modulating pluripotency around the peri-implantation stage^8^, enabling embryonic diapause^9^ and potentiating global transcriptional regulation^10^.

In previous reports we have described an elementary role for p38 mitogen-activated protein kinase (p38-MAPK) signalling on PrE specification in the mouse blastocyst inner cell mass (ICM)^12^, as well as a distinct function mitigating amino acid deprivation induced oxidative stress during blastocyst maturation^13^. Recently, we extended these investigations and described an early blastocyst functional role in regulating ribosomal RNA (rRNA) processing, translation and transcription required for correct mouse blastocyst expansion and PrE cell-fate specification and differentiation. Moreover, that this pathway appears largely independent of the quintessentially described FGF4-based PrE specification mechanisms and is at least partially upstream of the translation regulator mTOR^11^; itself independently implicated in PrE specific survival in late (E4.5) blastocysts^14^. p38-MAPK activity is also necessary for functional TE derivation prior to blastocyst formation, and its inhibition (p38-MAPKi) from the 16-cell stage (E3.0) specifically impairs fluid-filling of the blastocoel cavity by functionally impacting tight-junction proteins (TJP1), aquaporins (AQP3) and Na+/K+ pumps (such as ATP1); possibly as a result of p38-MAPKi induced translational and transcriptional deregulation^11,15,16^. Additionally, p38-MAPKi from the 8-cell stage (E2.5) is reported to cause developmental arrest, at the 8-16 cell stage transition, and is associated with defective embryo compaction and impaired filamentous actin formation^17^.

In our recent publication describing the role of p38-MAPK in protein translation regulation and PrE differentiation, we reported the results of a phosphoproteomics screen for differentially expressed phosphoproteins in early mouse blastocysts ±p38-MAPKi; aimed at identifying relevant potential p38-MAPK effectors/substrates^11^. This analysis identified the Myb-binding protein 1A (MYBBP1A), a known regulator of rRNA transcription and processing^18^ (shown by us to be defective in p38-MAPKi blastocysts^11^). We showed siRNA induced clonal downregulation of *Mybbp1a* was associated with cell-autonomous proliferation defects and a strong bias against PrE differentiation within mouse blastocyst ICM^11^. The HIV Tat-specific factor 1 homolog (HTATSF1) protein, itself implicated in genetic knockout studies as regulating peri-implantation stage blastocyst EPI pluripotency^8^, is a known functional interacting partner of MYBPP1A^8,19^. HTATSF1 also functionally associates with DEAD-box RNA helicase 21 (DDX21)^8,20^. Interestingly, although *Ddx21* was not specifically identified in our published ±p38-MAPKi early blastocyst screen^11^ (falling outside the prescribed filters), we had previously observed it as a differentially phosphorylated protein in our preliminary optimisation trials (supplementary table S1 – *derived from previously unpublished observations*). Therefore, we decided to revisit the, to date uncharacterised, functional role of *Ddx21* during mouse preimplantation development within the context of p38-MAPK activity.

*Ddx21* (initially termed RNA helicase II/Guα) is primarily, but not exclusively, a nucleolar-localised RNA helicase (with ATPase activity) shown to be a regulator of rRNA processing; downregulation of which causes decreased production of both 18S and 28S rRNA and accumulation of unprocessed 20S pre-rRNA transcripts^21,22^. It co-fractionates as a protein complex consisting of pre-60S ribosomal subunits and other rRNA processing proteins, such as Pescadillo homolog 1 (PES1), Eukaryotic translation initiation factor 4E-binding protein 2 (EIF4EBP2) and Guanine nucleotide-binding protein-like 3 (GNL3; also known as Nucleostemin/NS)^23^. Recent detailed molecular studies have reported diverse *Ddx21* roles that range from rRNA metabolism and regulation of RNA polymerase (pol) I and II mediated transcription to nucleotide stress response and c-Jun activity^20,24–27^. Indeed, *Ddx21* regulates RNA pol II mediated transcription of mRNA and non-coding RNA components of ribonucleoprotein complexes^20^. According to another recent report, *Ddx21* interacts with a specific long non-coding (lnc) RNA, termed the ‘small nucleolar RNA (snoRNA)-ended lncRNA that enhances pre-rRNA transcription’ or SLERT, to reduce its inhibitory affinity for the RNA pol I complex^26^. Regulation of *Ddx21* activity itself is under the control of post-translational modification, with CREB-binding protein (CBP) mediated acetylation and Sirtuin7 (SIRT7) mediated deacetylation respectively decreasing or increasing its helicase activity^28^. Whilst some, rRNA metabolism related mouse genes, for example *Mybbp1a*^18^, *Gnl3*^29^ and potentially *Htatsf1* (homozygous knockout embryonic stem cells demonstrate reduced pluripotency^8^), are associated with embryonic lethal genetic knockout phenotypes, no similar *Ddx21* specific studies are reported. However, the Mouse Genome Informatics (MGI) database does catalogue evidence that N-ethyl-N-nitrosourea (ENU) induced mutagenesis of the *Ddx21* gene is embryonically lethal (http://www.informatics.jax.org/marker/MGI:1860494). Ribosomopathies, a collective term for varied human congenital developmental disorders, are mostly associated with heterozygous mutations in factors involved in ribosome biogenesis^30,31^. Interestingly, experimental perturbation of disease-causing candidate genes, such as those identified in Treacher Collins syndrome (TCS), Diamond–Blackfan anaemia (DBA) and Shwachman–Diamond syndrome (SDS), have been reported to result in shifted localisation of DDX21 from nucleoli to the general nucleoplasm, with associated changes in DDX21 target chromatin interaction and rRNA processing^24^. Additionally, DDX21 can form a complex with the serine/threonine phosphoprotein phosphatase (PPP) family protein PP1^32^, that along with other family members are well characterised late M-phase/cytokinesis cell cycle regulators^33^.

Here, using pharmacological inhibition, siRNA mediated downregulation, confocal immuno-fluorescence microscopy and image analysis, we report our findings on the role of DDX21 during mouse preimplantation embryo development. We find basal levels of nuclear DDX21 protein expression until the 16-cell (E3.0) stage, when DDX21 levels begin to increase and become robustly expressed in the early (E3.5) blastocysts before becoming nucleolar by the late blastocyst stage. Moreover, that DDX21 protein expression is sensitive to p38-MAPKi during the period of blastocyst maturation (E3.5-E4.5), as evidenced by varying degrees of decreased nucleolar retention. Specific siRNA-mediated targeting of the *Ddx21* gene leads to efficient global mRNA knockdown and associated clonal cell-autonomous reductions in DDX21 protein expression that cause a 50% reduction in the volume of the late blastocyst. Intriguingly, such clonal *Ddx21* downregulation both cell autonomously (within the siRNA treated clone) and non-cell autonomously (affecting non-microinjected cells outside the *Ddx21* knockdown clone) impairs blastomere proliferation and results in severe defects in the numbers of EPI and PrE lineages by the late blastocyst stage; compared to equivalent stage non-specific control siRNA treated groups. These results complement our recent report on the general translational regulatory role of p38-MAPK during blastocyst development^11^ and identify DDX21 as a p38-MAPK effector with an apparently essential role in murine preimplantation embryonic development.

## Results

### DDX21 localisation shifts from nuclear to nucleolar post-cavitation during preimplantation development

We previously described an important role of p38-MAPK in coordinating an early blastocyst protein translation response necessary to promote PrE differentiation; identifying MYBPP1A, a known rRNA processing factor, as a p38-MAPK effector^11^. Whilst, there are mouse genetics-based evidences implicating some rRNA metabolism related genes (*e*.*g. Mybbp1a*^18^, *Gnl3*^29^ and *Htatsf1*^8^) to critical roles in early development, no similar data exists for the RNA helicase encoding *Ddx21* gene. Therefore, given the precedents for early developmental functions of similar rRNA-related genes, and the fact we have detected differential phosphorylation of DDX21 protein in mouse blastocysts after p38-MAPKi (supplementary table S1), we decided to assay *Ddx21* gene expression throughout the preimplantation developmental stages. Data from recently published transcriptomic studies report *Ddx21* mRNA expression starting to rise from a low steady state level at the 2-cell (E1.5) stage, peaking at the 8-cell stage and thereafter being maintained at a high level in all preimplantation embryonic cell lineages (supplementary Fig. S1)^34,35^. Immuno-fluorescent staining of DDX21 protein revealed expression at basal levels in the nuclei of 2-, 4-(E2.0) and 8-cell embryos and robust expression in a subset of 16-cell stage nuclei, that was found in all nuclei by the early through late blastocyst stages, correlating with formation of the blastocyst cavity (Fig. 1a). Interestingly, whereas in pre-cavitated embryos, DDX21 protein appeared to be either pan-nucleoplasmic or even adjacently localised to the nuclear membrane (Fig. 1b), post cavitation it was found to be exclusively nucleolar (Fig. 1d); suggesting a potential functional significance associated with the onset of blastocyst formation. Equally intriguing was the observation of similar differential DDX21 localisation behaviour regarding mitotic chromatin. DDX21 readily localised to condensed chromosomes prior to cavitation but appeared to be actively excluded from them post-cavitation (compare Figs. 1c & e); further suggesting functional adaptation of the role DDX21 as a response to blastocyst formation.

**Figure 1:**
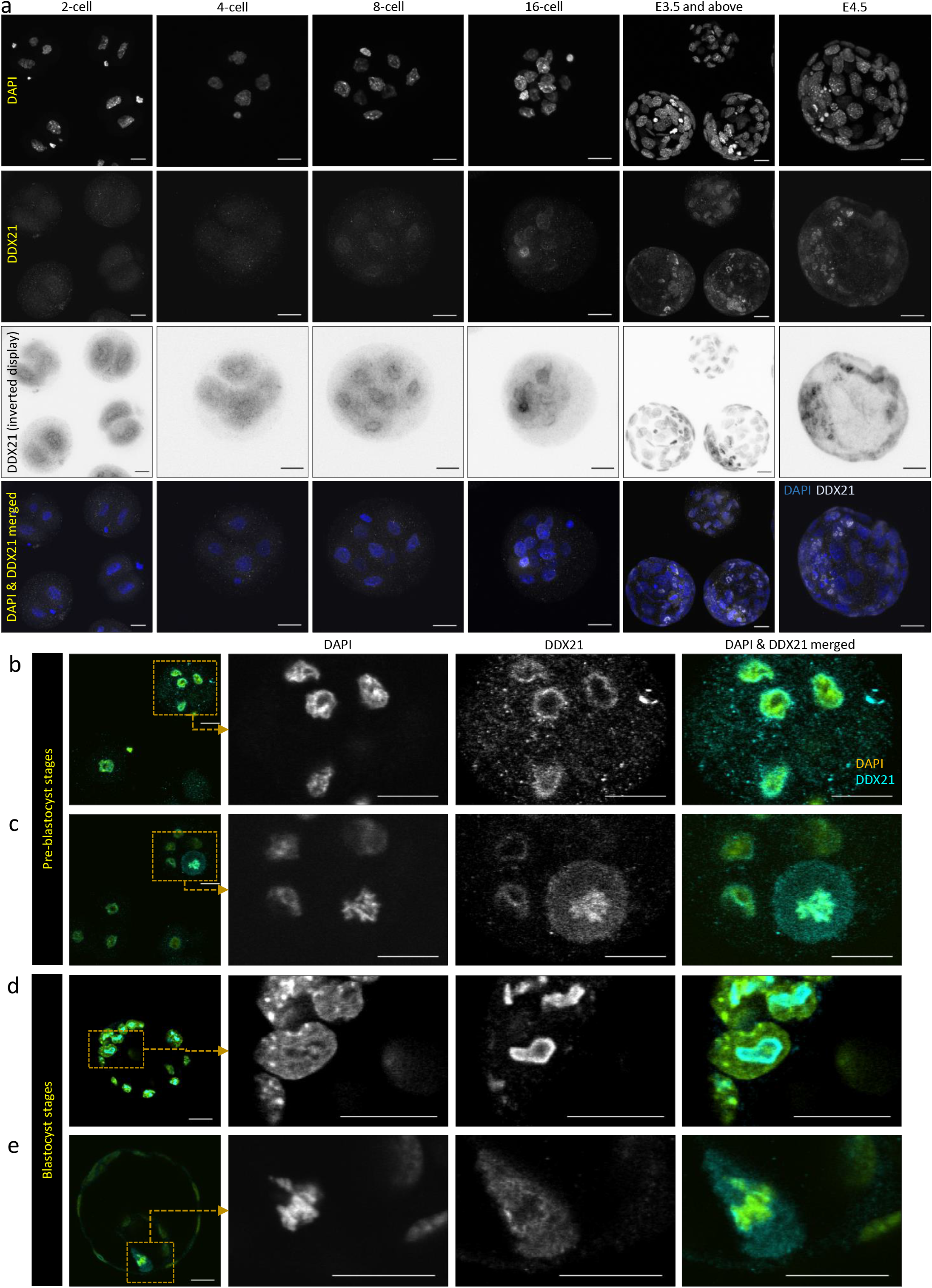
DDX21 protein expression and localisation in developing preimplantation mouse embryos. a) From left to right, Z-projection confocal micrographs of 2-cell, 4-cell, 8-cell, 16-cell, E3.5 and E4.5 embryos stained for the nucleus (DAPI), DDX21 (also displayed as an inverted grayscale image) and a merge of the two (DAPI pseudo-coloured blue, DDX21 in grayscale). b) to e) Single z-section confocal micrograph of an 8-cell stage embryo immuno-stained for DDX21 with magnified inlay showing (b) nucleoplasmic/nuclear membrane localisation in interphase cells and (c) condensed chromatin localisation in a mitotic cell. d) Single z-section confocal micrograph of a cavitated blastocyst-stage embryo immuno-stained for DDX21 with magnified image showing nucleolar localisation in interphase cells and (e) excluded localisation from condensed chromatin in a mitotic cell. DAPI and DDX2 signal is pseudo-coloured cyan and yellow in the merged images (panels b-e) and scale bar (panels a-e) = 20µm.

### DDX21 localisation is sensitive to p38-MAPK signalling during blastocyst maturation

Collectively, we have previously identified an early blastocyst developmental window of p38-MAPK activity (associated with protein translation regulation) required to support PrE differentiation^11^, obtained evidence DDX21 is a candidate p38-MAPK substrate (supplementary table S1) and revealed blastocyst specific nucleolar restricted expression of DDX21 protein. We therefore sought to examine DDX21 protein expression and localisation (together with that of the functionally related and rRNA processing factor Nucleostemin/NS, also known as Guanine nucleotide-binding protein-like 3/GNL3^23^; genetic nulls of which do not develop beyond E4.0^36,37^.) in maturing mouse blastocysts ±p38-MAPKi (E3.5-E4.5 -Fig. 2a). In control E4.5 stage blastocysts, we observed expression of DDX21 and GLN3 in both inner and outer cell nuclei, that in respect to nucleolar structures was co-localised (Fig. 2b & b’ insets). Nevertheless, respective protein expression quantitation revealed DDX21 to be more highly expressed in outer versus inner cell nuclei overall, with the inverse being true for GLN3 (compare Figs. 2 e & f, also alternatively expressed in supplementary Fig. S2). p38-MAPKi resulted in a clear reduction in both DDX21 and GNL3 protein expression that was evident in all blastocyst cells, comprising both inner and outer populations (Fig. 2c, d, e, f & supplementary Fig. S2). The percentage decrease upon p38-MAPKi in levels of GLN3 (56% decrease overall) was greater than DDX21 (30% decrease overall), nevertheless the values were statistically significant in both cases (Fig. 2c, d, e, f & supplementary Fig. S2). Apart from the changes in general DDX21 protein expression, we also note that p38-MAPKi was associated with a shift in DDX21 protein localisation. This was manifest in increased expression in the general nucleoplasm (Fig. 2c’ & d’) as opposed to exclusively nucleolar expression previously characterised in untreated (Fig. 1d) or control treated late blastocysts (Fig. 2b,b’). Interestingly, we did also observe the degree of p38-MAPKi dependent relocalisation of DDX21 protein away from the nucleolus was somewhat heterogeneous (compare Figs 2c & c’ with 2d & d’). This is possibly related to natural heterogeneity in the developmental timings of individual blastocysts at the point of administering p38-MAPKi. Notwithstanding, these data demonstrate the continued general expression of DDX21 and more profoundly GLN3 is sensitive to active p38-MAPK signalling during blastocyst maturation. Additionally, typical blastocyst stage nucleolar localisation of DDX21 protein is also regulated by active p38-MAPK. Collectively, these data lend support to the importance of p38-MAPK related regulation of translation, and particularly rRNA processing, as recently uncovered in early blastocyst maturation and eventual PrE differentiation^11^.

**Figure 2:**
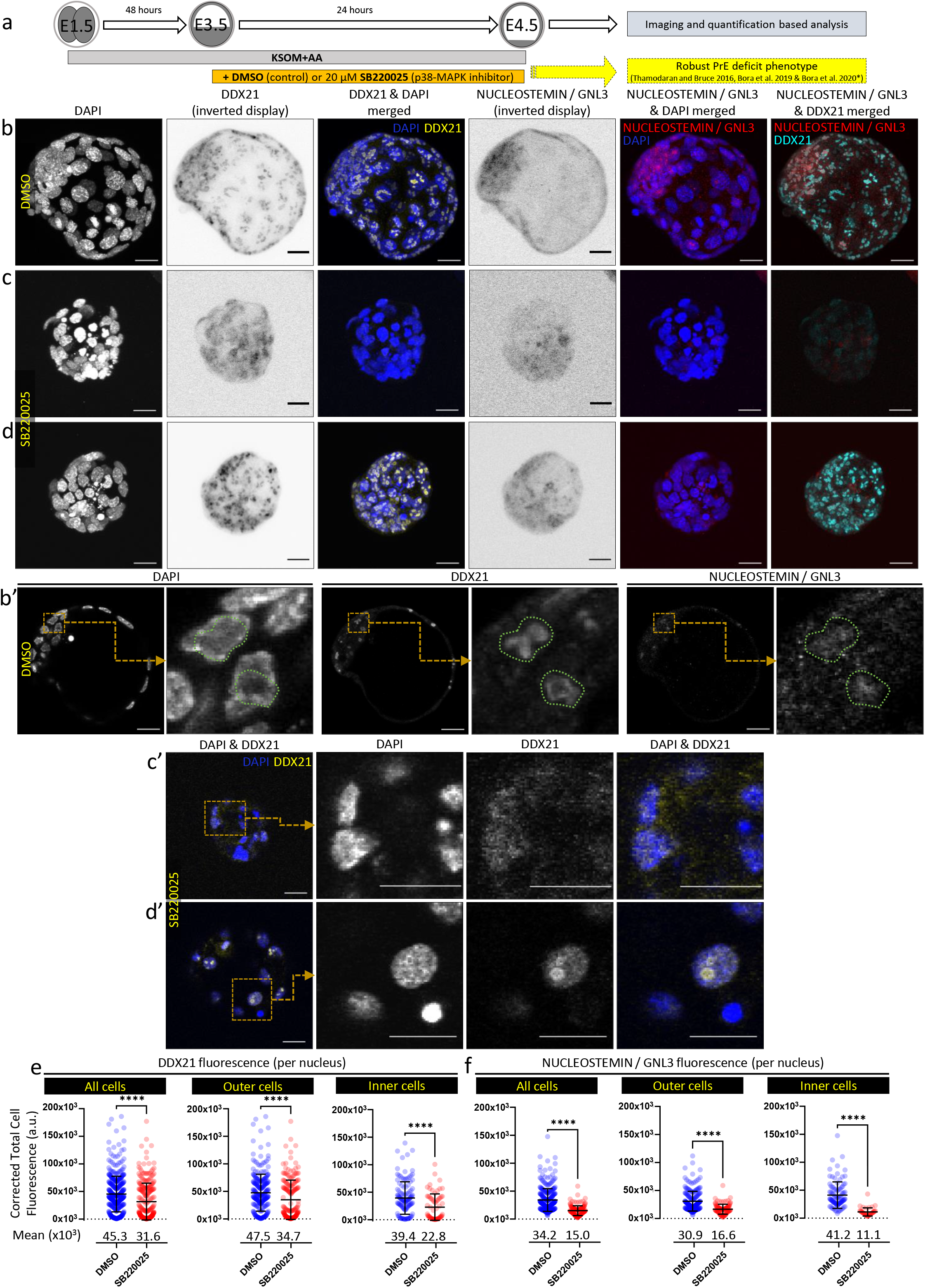
Effect of p38-MAPK inhibition on DDX21 and NUCLEOSTEMIN (GNL3) protein expression and localisation during blastocyst maturation. a) Scheme illustrating experimental protocol of p38-MAPK inhibition (p38-MAPKi) during blastocyst maturation (E3.5-4.5; previously reported by us to be a period during which p38- MAPK signalling is required for PrE specification and blastocyst maturation^11–13^). b) d) Confocal microscopy z-projections of (b) control (DMSO treated; n=7) and (c, d) p38-MAPKi (SB220025 treated; n=5) embryos immuno-stained for DDX21 and NUCLEOSTEMIN (GNL3) (both displayed as inverted grayscale images). In DAPI (blue) merged images, DDX21 and NUCLEOSTEMIN signal is pseudo-coloured yellow and red, respectively; in DDX21 and NUCLEOSTEMIN merges, respective cyan and red pseudo-colour pallets are used. b’) Single confocal z-sections, with magnified inlays, of DAPI, DDX21 & NUCLEOSTEMIN immuno-stained nuclei of control (DMSO treated) blastocysts. Green dotted enclosure demarcates nucleolar localisation of DDX21 and the region is superimposed on DAPI and NUCELOSTEMIN images highlighting co-localisation. c’) & d’) Magnified single confocal z-sections, with magnified inlays of DDX21 immuno-stained nuclei (co-stained with DAPI) of p38-MAPKi (SB220025 treated) blastocysts. In merged images, DAPI and DDX21 signals are pseudo-coloured blue and yellow, respectively. Scale bar (panels b-d) = 20μm. e) & f) Scatter plot quantifications of per cell nuclei Corrected Total Cell Fluorescence (CTCF) of e) DDX21 and f) NUCLEOSTEMIN (GNL3) immuno-staining in control (DMSO treated) and p38-MAPKi (SB220025 treated) embryos, as expressed for all cells, outer cells and inner cells. DDX21: DMSO from n=16 embryos (outer cells=271, inner cells=122) & SB220025 from n=15 embryos (outer cells=190, inner cells=67) and NUCLEOSTEMIN: DMSO from n=7 embryos (outer cells=205, inner cells=97) & SB220025 from n=5 embryos (outer cells=100, inner cells=40). Collated individual nuclei CTCF quantifications for (e) and (f) are provided in supplementary table S4. (Alternative comparative representations of these data are provided in supplementary Fig. S2)

### Clonal downregulation of *Ddx21* causes reduced embryo cell number and blastocyst expansion

Given the revealed p38-MAPKi sensitivity of blastocyst DDX21 nucleolar localisation, we speculated what the effect of targeted dysregulation of the *Ddx21* gene expression would be on mouse preimplantation embryo development and, given our previously described reports^11–13^, on blastocyst ICM cell fate derivation and specifically PrE differentiation. Accordingly, we devised a siRNA microinjection mediated scheme to downregulate *Ddx21* transcript levels (schematic described in Fig. 3a). To assess *Ddx21* specific siRNA efficiency, we microinjected both blastomeres of 2-cell stage embryos and quantified normalised *Ddx21* transcript levels at the late blastocyst stage. Compared to non-targeting control (NTC siRNA) microinjected embryos, at the equivalent stage, *Ddx21* specific siRNA elicited 42% decreased *Ddx21* mRNA expression (Fig. 3b). We next, created fluorescently marked embryonic clones by co-microinjecting one cell at the 2-cell stage with siRNA (either NTC or *Ddx21* specific) and mRNA encoding histone H2B-RFP reporter; thus, creating a clone initially comprising 50% of the embryo. Microinjected embryos were then cultured to the late blastocyst stage, fixed, immuno-fluorescently stained for DDX21 (and the outer trophectoderm marker protein CDX2 ^38^– to distinguish inner and outer cell populations) and the number of cells in the fluorescently marked siRNA microinjected clones, plus non-microinjected sister clones, and their spatial localisation within the embryo (outer or inner) recorded.

**Figure 3:**
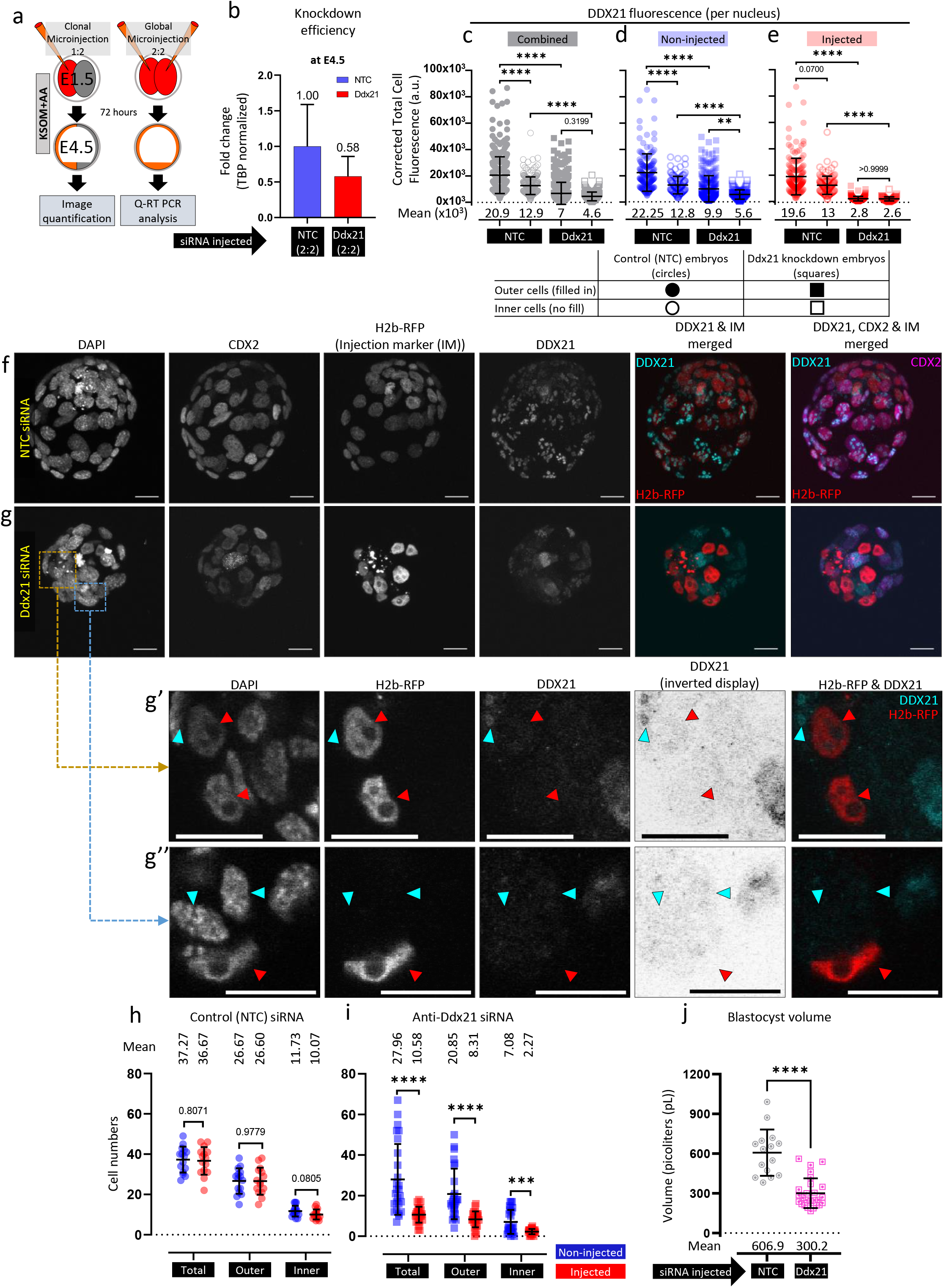
Clonal knockdown of *Ddx21* and effect on late-blastocyst morphology and cell numbers. a) Experimental design to determine the efficiency of siRNA mediated *Ddx21* gene mRNA knockdown in microinjected embryos cultured to the equivalent late blastocyst (E4.5) stage (right panel) and to assay the contribution of marked *Ddx21* knockdown clones to late blastocyst cell lineages (left panel) b) qRT-PCR derived relative *Ddx21* transcript levels (normalised to *Tbp* mRNA levels) between embryos injected with non-targeting control (NTC) siRNA and siRNA specific for *Ddx21* mRNA at E4.5. c)-e) Scatter plot quantifications of per cell CTCF values for DDX21 expression in control (NTC siRNA injected n=15 embryos; non-injected cells outer n=151 & inner n=108, injected cells outer n=153 & inner n=112) and *Ddx21* downregulated (*Ddx21* siRNA injected n=26; non-injected cells outer n=225 & inner n=125, injected cells outer n=157 & inner n=59) embryos,comparing c) all inner and outer cells, between NTC and *Ddx21* siRNA injected embryos, d) only non-injected cells between inner and outer cells for both sets of embryos, and e) only injected cells between inner and outer cells for both sets of embryos (Collated individual nuclei CTCF quantifications for (c), (d) and (e) are provided in supplementary table S5). f) & g) Confocal micrograph z-projections of representative late (E4.5) stage blastocysts initially microinjected (in one blastomere at the 2-cell stage) with (f) NTC siRNA (n=15) and (g) *Ddx21* siRNA (n=26), plus recombinant *H2b-RFP* fluorescent reporter mRNA (identifying the clonal progeny of the injected cell). Individual DAPI (grayscale pan-nuclear stain; total number of cells), DDX21 (grayscale) and CDX2 (grayscale; TE cells) channel micrographs, plus merged DDX21 (cyan), CDX2 (magenta) and H2B-RFP (red; microinjected clone) images are shown. g’) & g’’) Magnified single z-section confocal micrograph of clonal *Ddx21* downregulated embryo with DDX21 immuno-stained nuclei representative for both autonomous and non-autonomous effects (and DAPI counterstain). In merged image H2B-RFP and DDX21 signal is pseudo-coloured red and cyan, respectively. Red arrowheads mark progeny of *Ddx21* siRNA microinjected cells (discernible by H2B-RFP fluorescence – detailing cell autonomous DDX21 protein knockdown) and cyan arrowhead marks non-microinjected cells (detailing in g’) continued but reduced DDX21 expression, or g’’) non-autonomous DDX21 protein knockdown). Scale bar (panels f-g’’) = 20μm. h) & i) Scatter plot quantification of cell numbers in (h) NTC siRNA and (i) *Ddx21* siRNA microinjected embryos, categorised into total, inner and outer cell populations, based on combined CDX2 expression and DAPI fluorescence. Collated individual embryo cell number quantifications are provided in supplementary table S7. (Alternative comparative representations of these data are provided in supplementary Fig. S3) j) Quantification of blastocyst volume (pL). Collated individual embryo volume quantifications are provided in supplementary table S6.

As shown in the confocal micrographs (Fig. 3f-g’’), microinjected cell clones were clearly distinguishable by expression of the histone H2B-RFP reporter in both NTC and *Ddx21*-specific siRNA microinjection groups. However, whereas DDX21 protein was expressed in all cell nuclei (specifically nucleoli) in the NTC control group, it was only readily detected in cell nuclei derived from the non-microinjected clone in the *Ddx21* targeted group (although not always restricted to nucleoli) and appeared highly, if not completely, reduced in the marked microinjected clone (Fig. 3g, g’ & g’’). Indeed, normalized quantitation of DDX21 immuno-fluorescence confirmed a highly significant reduction in overall average expression per cell (encompassing both microinjected and non-microinjected cell clones) in the *Ddx21* siRNA treatment group versus the NTC control (Fig. 3c-e & alternatively presented in supplementary Fig. S3a-c). This was true of both outer and inner cell populations (note, DDX21 expression was higher in outer versus inner cells in the NTC siRNA controls and was therefore consistent with observations made on similar late blastocysts treated with vehicle control DMSO – Fig. 2 & Fig. S2). The *Ddx21* siRNA induced reduction in DDX21 protein expression was most robust in the marked microinjected clone in both inner and outer cells (*i*.*e*. when compared to either the non-microinjected sister clones of the same embryos or the equivalent microinjected clone of the control NTC siRNA treatment group (Fig. 3e)); thus, confirming the anticipated cell autonomous effect of the microinjected *Ddx21* siRNA. However, it was also observed that clonal *Ddx21* knockdown also affected DDX21 protein expression in the non-microinjected clone, as it was also significantly reduced (in both inner and outer cells) in *Ddx21* versus NTC siRNA treatment groups (Fig. 3d and g’’); confirming additional DDX21 related non-cell autonomous effects (that nevertheless did not affect every cell within that clone (Fig. 3d and g’). Notably, the microinjection of NTC siRNA had no statistically significant effect on DDX21 protein levels in either inner or outer cells and confirmed the absence of bias that could be attributable to the microinjection procedure. Hence, microinjection of *Ddx21* siRNA in one blastomere at the 2-cell stage causes highly efficient DDX21 protein knockdown within the clonal progeny of the microinjected cell but also results in reduced DDX21 expression in the accompanying non-microinjected sister clone, that is not restricted to the nucleolus by the late blastocyst stage.

Inspection of the embryos derived from the clonal NTC and *Ddx21* siRNA treatment groups also revealed profound morphological differences. Whereas, the NTC group comprised embryos with typical late blastocyst morphology (*i*.*e*. a mature CDX2 positive outer trophectoderm layer and large blastocyst cavity and ICM), the *Ddx21* siRNA microinjected embryos were typically smaller and appeared to comprise much fewer cells (although CDX2 protein expression was appropriately confined to outer cells only – compare Fig. 3f &g). Indeed, measuring the volume of the derived blastocysts, we observed a more than 50% reduction in *Ddx21* clonally downregulated embryos compared to control embryos (Fig. 3j). This result resonates with our recent report in which we revealed a similar blastocyst expansion defect, that is associated with significantly impaired PrE differentiation, upon p38-MAPKi (E3.5-E4.5)^11^ and the fact we have identified DDX21 as a potential p38-MAPK effector (supplementary table S1). An analysis of the average number, relative spatial positioning and clonal origin of cells showed no significant differences between the NTC siRNA microinjected clone and its non-microinjected counterpart. However, in the *Ddx21* siRNA group only 27.45% of overall cell numbers originated from the microinjected blastomere (compared to 49.6% for NTC microinjected embryos), with comparative cell number deficits in the microinjected (versus non-microinjected) clone being highly significant in both outer and inner cell populations (on average 8.31 versus 20.85 and 2.27 versus 7.08, respectively) (Fig. 3h & I and supplementary Fig. S3d). The reduced contribution was particularly stark for inner cells, averaging at 24.3% of the total ICM (compared to 46.2% in controls), that in absolute numbers only represents an average of 2.27 inner cells. In addition to such *Ddx21* siRNA microinjected clone specific cell deficits, we also observed significantly reduced numbers of outer and inner cells in the non-microinjected clone (when compared to the equivalent clone in the control NTC siRNA treatment group – on average 20.85 versus 26.67 and 7.08 versus 11.73, respectively) (Fig. 3h & i). These data confirm both cell autonomous and non-autonomous effects of clonal *Ddx21* knockdown in developing mouse embryos and agree with the atypical non-nucleolar localisation of DDX21 in the non-microinjected clone (see above). Thus, clonal downregulation of *Ddx21* results in a global effect on embryonic development, effecting cell proliferation and blastocyst volume.

### *Ddx21* downregulation results in defective blastocyst cell-fate specification

Having previously confirmed a role for p38-MAPK during blastocyst maturation and PrE lineage differentiation^11–13^ plus the effects of p38-MAPKi on DDX21 protein localisation (Fig. 2) and the remarkably reduced contribution of *Ddx21* siRNA injected blastomeres towards inner cells (Fig. 3), we wanted to assay the effect of clonal *Ddx21* knockdown on late blastocyst (E4.5) ICM cell fate. Accordingly, we repeated our clonal NTC/*Ddx21* siRNA 2-cell stage microinjection experiments (as described in Fig. 3a) and assayed the expression of NANOG and GATA4 protein as markers of the EPI and PrE. Consistently, we again observed robust and significant decreases in overall, outer and inner cell numbers in *Ddx21* siRNA microinjected embryos, that was most evident in the microinjected clone but also present in the non-microinjected clone (supplementary Fig. S4). We also observed significant and marked reductions in the overall number of ICM cells expressing either NANOG or GATA4, in both microinjected and non-microinjected clones, in *Ddx21* siRNA treated embryos (Fig. 4a, b & c). Whilst these reductions can reflect overall reduced cell number, it is notable the effect was stronger for GATA4 expressing PrE versus NANOG positive EPI-like cells, with an overall average of 1.0 GATA4 expressing cell in *Ddx21* downregulated embryos compared to 7.44 in control NTC siRNA embryos (Fig. 4c (ii) – the equivalent NANOG expressing EPI cell numbers being 3.47 and 9.0, respectively – Fig. 5c (i)). Moreover, among the *Ddx21* downregulated embryos, we observed an average of 0.93 GATA4 expressing cells originating from the non-injected blastomere and only 0.07 originating from the injected one (Fig. 4c (ii) -the equivalent NANOG expressing EPI cell numbers being 1.93 and 1.53, respectively -Fig. 4c (i)). Hence, despite the reduced number of overall, and specifically ICM, cells associated with clonal *Ddx21* downregulation, it is the derivation of GATA4 expressing PrE lineage that is more markedly impaired/blocked by the late blastocyst stage. This is further revealed by the significantly reduced ratio of GATA4 positive PrE cells versus total ICM cell number, in both the microinjected and non-microinjected clones, of *Ddx21* siRNA treated embryos (Fig. 4c (iii)); a trend not observed in the calculated ratio of NANOG positive EPI-like cells in the same embryos (Fig. S4 (iv)). These data support a role for DDX21, potentially regulated by p38-MAPK activity, in facilitating PrE differentiation in the mouse blastocyst.

**Figure 4:**
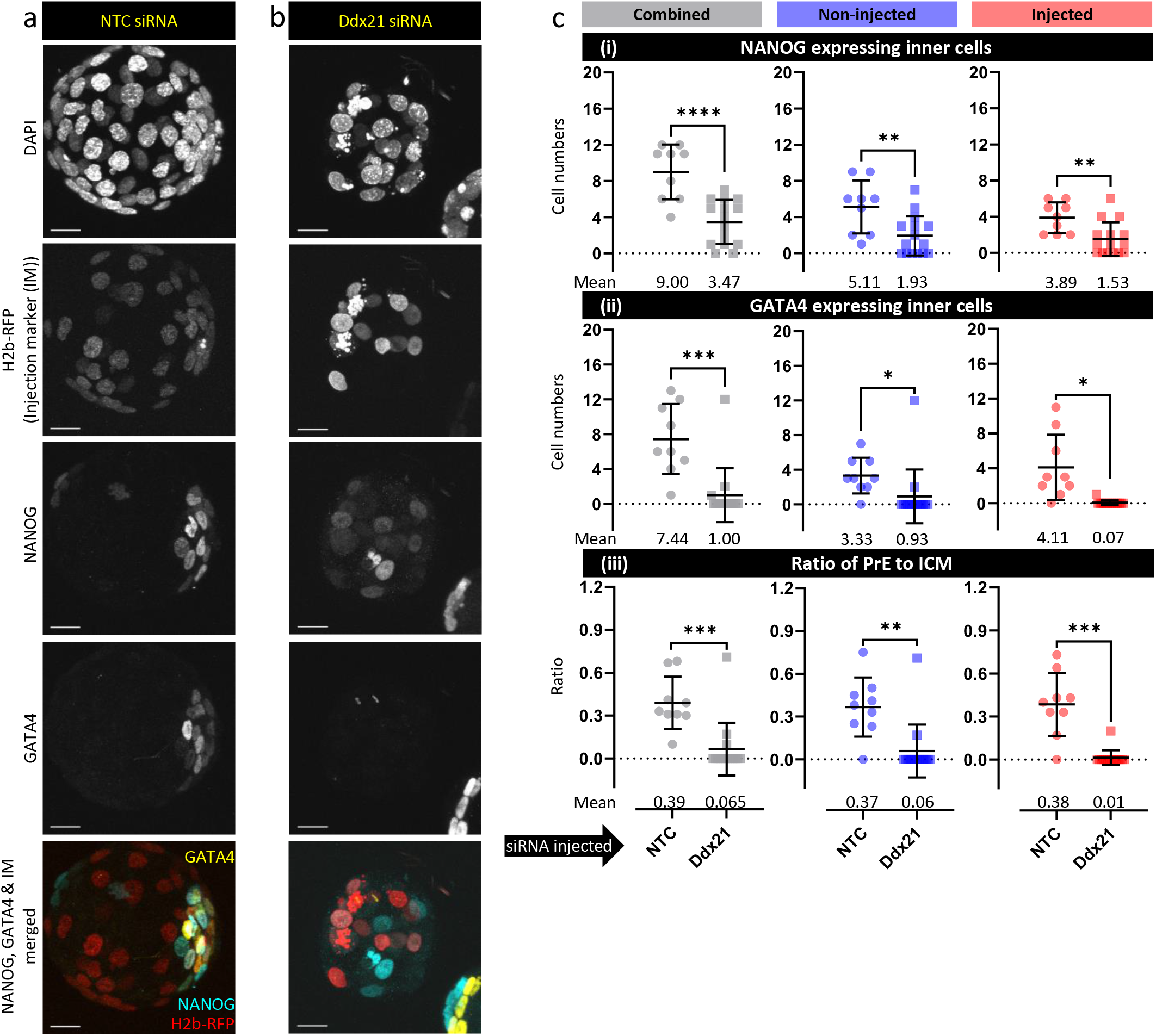
Effect of *Ddx21* knockdown on Epiblast (EPI) and Primitive Endoderm (PrE) lineage specification at E4.5. a) & b) Confocal microscopy z-series projections of late blastocyst (E4.5) stage embryos derived from clonal microinjections (one blastomere at the 2-cell stage) of a) control NTC siRNA (n=9) and b) *Ddx21* siRNA (n=15) embryos immuno-stained for EPI (NANOG) and PrE (GATA4) lineage markers; note H2B-RFP distinguished microinjected and non-microinjected cell clones (experimental design in Fig. 3a). In merged images, NANOG, GATA4 and H2B-RFP signals are pseudo-coloured cyan, yellow and red respectively. Scale bar = 20μm. c) Scatter plot quantification of (i) EPI cell numbers (only expressing NANOG), (ii) PrE cell numbers (only expressing GATA4) and (iii) the PrE to total ICM ratio, of clonal NTC siRNA and *Ddx21* siRNA microinjected injected embryos, as observed in either the microinjected or non-injected clones and both clones combined. Collated individual embryo cell number quantifications are provided in supplementary table S8. (Alternative comparative representations of these data are provided in supplementary Fig. S4)

## Discussion

The functional role of p38-MAPKs in preimplantation mouse embryos is a developing story and work from a few labs, including ours, have only just started to uncover the many facets of early development that this signalling pathway touches. In the earlier developmental stages, p38-MAPK activity is associated with formation of filamentous actin^17^ and embryo compaction and subsequent appropriate functioning of the TE to enable expansion of the blastocoel cavity^15,16^. Our own p38-MAPK research has primarily focussed on the post E3.5 cavitated stages, thus specifically addressing blastocyst maturation and ICM specification towards EPI and PrE lineages^11–13^. In our recent work, we have identified a protein translation associated regulatory role for p38-MAPK that underpins PrE cell fate differentiation. This role appears to be functionally upstream of the mTOR pathway and largely independent of the classically described FGF4-ERK mediated mechanisms of PrE specification^11^. Whereas the upstream regulators/activators of this p38-MAPK pathway are the subject of informed speculation, potentially involving FGFR2, FGFR3 and/or PDGFRa^11^, the downstream effectors can be many and encompass various mechanistic ontologies (reviewed here^39^). Hence the identification of known factors involved in rRNA metabolism and ribosome biogenesis (for example, MYBBP1A^11^ and DDX21 described here) as being sensitive to p38-MAPK signalling and essential in appropriate preimplantation embryonic development, further reinforces the emerging importance of the broader hypotheses of germane regulation of protein translation.

To the best of our knowledge, we report the first observations of DDX21 protein expression and localisation during mouse preimplantation embryonic development (Fig. 1a). Expression of the *Ddx21* gene is reported to start around the 2-4 cell stage and peaks at the 8-cell stage in mouse embryos, subsequently remaining high across all cellular lineages; with close to nil expression in the maturing oocyte (supplementary Fig. S1)^34,35^. The protein appears to be expressed only after ZGA onset, and is thus not of maternal origin, becoming robustly observable in some 16-cell stage nuclei. Around blastocyst formation, all cells express DDX21 protein that then exhibits a typical nucleolar localisation as the blastocyst matures to the peri-implantation stage (Fig. 1a & d). Thus, the protein expression timeline closely follows that of the gene transcript. Moreover, the fact *Ddx21* knockdown clones only contribute an average of 11.69 cells to the late blastocyst, compared to 36.67 cells from the NTC siRNA microinjected blastomeres (Fig. 3h & i)), strongly suggests DDX21 expression is ordinarily necessary for transition beyond the 32-cell stage and in turn blastocyst formation. This is reinforced by the observation that such cell autonomous clonal cell number effects are also accompanied by non-cell autonomous effects within the non-microinjected clone; implying the existence of inter-blastomere proliferative signals that are impinged upon by functional DDX21. Moreover, the existence of such DDX21 related non-cell autonomous effects is also suggested by the fact DDX21 protein expression in the non-microinjected clone of *Ddx21* siRNA microinjected embryos does not always exhibit nucleoli restricted expression typical of the blastocyst stages. Nucleoli are sub-nuclear structures closely associated with ribosome biogenesis that gradually transit through extensive compositional, functional and structural transformations between oogenesis and the embryonic morula stage (reviewed here^40,41^). Briefly, as developmentally competent oocytes slowly initiate inactivation of RNA pol I and II, their nucleoli gradually become denser and form atypical structures termed nucleolus-like-bodies (NLB); a process that is accompanied by reducing levels of component nucleolar proteins and correlates with paused ribosome biogenesis. After fertilisation, further atypical nucleolar bodies, termed nucleolus-precursor-bodies (NPB), emerge but are then replaced by somatic cell-type nucleoli after the morula to blastocyst transition. The functional significance of these atypical nucleolar bodies during early development is itself an interesting topic, with some reports suggesting their requirement during early ZGA and as gene regulatory structures in later stages^41^. However, it is interesting that our observations of exclusive nucleolar restricted DDX21 protein expression is concomitant with the emergence of somatic cell-type nucleoli during the earliest stages of blastocyst formation. Moreover, that such exclusive nucleolar expression is dependent on active p38-MAPK during blastocyst maturation, a time in which p38-MAPK mediated regulation of protein translation is required for PrE differentiation^11^. Thus, it is possible DDX21, potentially acting downstream of p38-MAPK regulation, is involved in the remodelling of early embryonic cell nucleoli, in order to sustain the morula to blastocyst transition and subsequent ICM maturation and PrE formation.

Related to the subject of DDX21 protein localisation, the observed differential association of DDX21 with condensed mitotic chromosomes as a function of developmental stage is intriguing (*i*.*e*. in association in pre-blastocyst stages and excluded in the blastocyst -Fig. 1c and e). There is a report suggesting DDX21 forms complexes with members of PPP family of phosphatases^32^ and indeed PP1 and other members of PPP family are key cell cycle regulators^33^. Thus, this association may explain the observed DDX21 localisations in relation to mitotic chromatin but the stage specific differential nature of such localisation also implies any potential DDX21-PPP interaction is likely to be dynamically regulated during preimplantation development. Interestingly, it is during blastocyst formation that mouse embryo mitoses revert to centrosomal control, after previously being acentrosomally regulated since fertilisation (a murine specific characteristic amongst studied mammalian species^42^).

Post-translational modification by acetylation is reported to reduced DDX21 activity^28^. Here we report reduced phosphorylation of DDX21 at serine-218 (S218) in mouse blastocysts upon p38-MAPKi (supplementary table S1). According to the phosphosite.org database (an archive of all confirmed and predicted phosphorylation events in a host of model organisms^43^) this site is mouse specific and not conserved in either humans or rats; however, it does conform with the consensus [S/T]P dipeptide phosphorylation motif typical of the proline-directed mitogen-activated-kinase superfamily^44^. Direct experimental evidence of p38-MAPK phosphorylation mediated functional and/or locational regulation of DDX21 is lacking. However, based on our current observations this potential mechanism remains a strong possibility; as exemplified by the shift to exclusively nucleolar DDX21 localisation upon blastocyst formation, that is itself sensitive to p38-MAPKi (Fig. 1 and Fig. 2). Such shifts in DDX21 localisation are also observed in genetic models of congenital ribosomopathies, such as TCS, DBA and SDS, and are interestingly present in the differentiated derivative of ES cells harbouring the specific genetic mutations but not the originating pluripotent parental ES cell lines themselves^24^. Thus, the p38-MAPKi phenotypes we observe during blastocyst maturation are, at least in the context of DDX21 localisation, similar to those translation defects underpinning described ribosomopathies. Moreover, coupled with our recent report of reduced translation and rRNA processing in blastocysts cultured under p38-MAPKi conditions^11^, such ribosomopathy based insights^24^ may be able to contribute to an expanded picture of the role of p38-MAPK signalling during blastocyst maturation and specifically ICM cell lineage and PrE derivation.

Our analysis of the contribution of *Ddx21* downregulated cell clones to late blastocyst (E4.5) ICM lineages is ostensibly an extension of our previous observations obtained under p38-MAPKi culture conditions^11–13^, and is similarly characterised by a reduction in the number of GATA4 expressing PrE cells (Fig. 4). However, it is the severity of this deficit that distinguishes such *Ddx21* knockdown phenotypes, as almost no GATA4 expressing PrE cells were found to originate from *Ddx21* siRNA microinjected clone and moreover such GATA4 deficits were also non-cell autonomously observed in the non-microinjected sister clones (Fig. 4c (ii)). We similarly observed highly significant reductions in NANOG expressing EPI-like cells and outer TE cells in *Ddx21* siRNA microinjected embryos (Fig. 4c (i) and supplementary Fig. S4), although, compared to the almost complete block in PrE derivation, these lineages were less severely affected. In contrast, the p38-MAPKi (E3.5-E4.5) derived phenotype is more specifically centred on PrE cell differentiation (characterised by an uncommitted state of NANOG and GATA6 co-expression), with only limited and insignificant decreases in the quantity of other late blastocyst cell lineages^11–13^. Hence, these data suggest DDX21 likely plays a more fundamental role in regulating blastomere proliferation at the morula to blastocyst transition (supported by the increasing DDX21 protein expression data -Fig. 1) and developmentally precedes the previously studied period of p38-MAPKi during blastocyst maturation (E3.5-E4.5)^11–13^. Interestingly, it is not possible to study ultimate ICM cell fate in embryos exposed to p38-MAPKi before E3.5 (and the onset of cavitation and irreversible TE specification^45^) as embryos arrest development with around 32-cells and fail to cavitate^12^. This confirms the morula-blastocyst transition as acutely sensitive to p38-MAPKi and developmentally coincides with the observed apparent proliferative block of *Ddx21* knockdown embryos (*i*.*e*. observed as more severe in the microinjected clone versus non-cell autonomous effects in the non-microinjected clone -Fig. 3 and Fig.4) and the observed transition of DDX21 protein expression to an exclusively nucleolar localisation (Fig. 1). Clonal *Ddx21* knockdown is still associated with cavity formation, albeit impaired in comparison to controls (Fig. 3j). This suggests that at least part of the pre-E3.5 administered p38-MAPKi 32-cell arrested development phenotype could have a DDX21 related root. In relation to p38-MAPKi phenotypes during blastocyst maturation (E3.5-E4.5), it is likely any effect of p38-MAPK mediated regulation of DDX21 (*e*.*g*. potential direct phosphorylation leading to the observed loss of exclusively nucleolar expression – Fig. 2) will probably not be as drastic as efficient downregulation of the *Ddx21* derived mRNA and protein caused by RNAi. This leaves open the possibility active p38-MAPK regulation of DDX21 function is a contributing factor to our previously observed p38-MAPK role in protein translation, mouse blastocyst ICM maturation and PrE differentiation^11^. Much like our observations with *Ddx21* (this manuscript) and *Mybbp1a*^*11*^ clonal downregulation in preimplantation embryos, *Mybbp1a*^*18*^ and *Gnl3* (Nucleostemin)^29^ homozygous gene knockouts also result defective development by the late blastocyst stage. These collective observations are bringing into focus the indispensable and potentially non-redundant role of various ribosome biogenesis related genes during the preimplantation stages of mouse development and the formation of the blastocyst ICM lineages in particular. Moreover, they also highlight the importance of the underpinning regulatory signalling mechanisms, such as the p38-MAPK pathway, responsible for their proper functioning.

## Methods

### Mouse lines and embryo culture

All mouse related experimental procedures (*i*.*e*. collecting preimplantation stage embryos for further study) complied with ‘ARRIVE’ guidelines and were carried out in accordance with EU directive 2010/63/EU (for animal experiments). Superovulation and strain mating regimen to produce experimental embryos; F1 hybrid (C57Bl6 x CBA/W) females injected sub-peritoneally with 7.5 IU pregnant mare serum gonadotrophin/PMSG (Folligon, MSD Animal Health, Boxmeer, Netherlands) and 48 hours later with 7.5 IU recombinant human chorionic gonadotrophin/hCG (Sigma-Aldrich, St. Louis, Missouri, USA; Cat. No. CG10), followed by overnight F1 male mating (successful mating confirmed by presence of vaginal plugs). E1.5 (2-cell) stage embryos were isolated from oviducts in M2 media (pre-warmed at 37°C for at least 2-3 hours) and thereafter cultured in KSOM (Sigma-Aldrich, EmbryoMax KSOM Mouse Embryo Media; Cat. No. MR-020P-5F -pre-warmed and equilibrated in 5% CO_2_ and 37°C) with amino acid (AA) supplementation; Gibco MEM Non-Essential Amino Acids Solution (100X) (Thermo Fisher Scientific, Paisley, Scotland; Cat. No. 11140035) and Gibco MEM Amino Acids Solution (50X) (Thermo Fisher Scientific; Cat. No. 11130036) were used to a working concentration of 0.5X. Embryos were cultured in micro-drops prepared in 35mm tissue culture dishes covered with light mineral oil (Irvine Scientific, Santa Ana, California, USA; Cat. No. 9305), in 5% CO_2_ incubators maintained at 37°C until the appropriate stage. Pharmacological manipulations were performed by addition of chemical agents dissolved in dimethyl sulfoxide (Sigma-Aldrich; Cat. No. D4540) to KSOM+AA and equivalent volumes of DMSO were used as vehicle controls. Please note, the p38-MAPK (SB220025 – 20μM) inhibitor concentration used was derived by empirical titrations previously employed by ourselves^12^ and literature based precedents^15,46^. All KSOM based culture media, with or without additional chemicals (AAs, pharmacological agents), were pre-warmed and equilibrated in 5% CO_2_ and 37°C for at least 3-4 hours prior to embryo transfer.

### Sample collection for mass spectrometric analysis of differential (phospho)proteome

2-cell (E1.5) stage embryos were cultured in normal KSOM+AA conditions until E3.5 +2h and thereafter 100 embryos each were moved to control or p38-MAPKi conditions and cultured for another 7 hours (E3.5 +9h). The embryos were then washed through pre-warmed (37°C) Hank’s balanced salt solution (HBSS, Sigma-Aldrich; Cat. No. H9269) and lysed by moving to a 1.5ml centrifuge tube containing about 15µl of SDT-lysis buffer (4% (w/v) SDS, 100 mM Tris-HCl pH 7.6, 0.1 M DTT). Cell lysis was performed by incubating the tubes in a 95°C thermoblock for 12 minutes, brief centrifugation at 750 rpm, cooling to room temperature and storage at -80°C.

### Samples preparation for Liquid Chromatography–Mass Spectrometry (LC-MS) analysis

Individual protein solutions were processed and analysed as described previously^11^. Briefly, protein lysates were processed by the filter-aided sample preparation (FASP) method. FASP eluates were used for phophopeptides enrichment using High-Select TiO2 Phosphopeptide Enrichment Kit (Thermo Fisher Scientific, Waltham, Massachusetts, USA). LC-MS/MS analyses of peptide mixtures (not enriched and enriched in phosphopeptides using TiO2 enrichment kit) were performed using a RSLCnano system connected to Orbitrap Fusion Lumos mass spectrometer (Thermo Fisher Scientific) as previously specified^11^. Analysis of the mass spectrometric RAW data files was performed using the MaxQuant software (version 1.6.1.0) using default settings. MS/MS ion searches were executed against modified cRAP database (based on http://www.thegpm.org/crap, 111 protein sequences) containing protein contaminants like keratin, trypsin *etc*., and UniProtKB protein database for *Mus musculus* (ftp://ftp.uniprot.org/pub/databases/uniprot/current_release/knowledgebase/reference_proteomes/Eukaryota/UP000000589_10090.fasta.gz; version from 20.6.2018, number of protein sequences: 22,297). Other conditions were the same as previously described^11^. Protein intensities reported in proteinGroups.txt file and evidence intensities reported in evidence.txt file (output of MaxQuant program) were further processed using the software container environment (https://github.com/OmicsWorkflows), version 3.5.3c. Processing workflow is available upon request. Protein and phosphopeptide candidates were selected based on the following criteria; Differential phospho-peptide candidates (exhibiting ≥1.5 fold differential abundance; note the cut-off was applied to acknowledge the potential sensitivity threshold limitations involved using such scare starting material) were selected based on the criteria described previously^11^.

### Embryo manipulation by microinjections

Single (for immuno-fluorescence confocal microscopy) or double (for qRT-PCR) blastomere microinjections of 2-cell (E1.5) stage embryos were performed using the FemtoJet 4i (Eppendorf, Hamburg, Germany; Cat. No. 5252000013) micro-injector, mechanical micro-manipulators (Leica, Wetzlar, Germany; Cat. No. ST0036714) and CellTram Vario (Eppendorf; Cat. No. 5176000033) pneumatic handler, under a negative capacitance enabled current controlled by an Electro 705 Electrometer (WPI, Sarasota, Florida, USA; Cat. No. SYS-705) and on the stage of an Olympus IX71 inverted fluorescence microscope. Embryos were pneumatically handled and immobilised for microinjection using a borosilicate glass capillary holder (without filament -Harvard Apparatus, Holliston, Massachusetts, USA; Cat. No. 30-0017). Micro-injectors were connected to needles prepared from filamented borosilicate glass capillaries (Harvard Apparatus; Cat. No. 30-0038) using a Narishige PC-10 capillary glass needle puller (Narishige Scientific Instrument Lab., Tokyo, Japan). siRNAs were co- microinjected at 10µM concentrations, with 50ng/µl *H2b-RFP in vitro* transcribed (mMESSAGE mMACHINE T3 IVT kit, Thermo Fisher Scientific; Cat. No. AM1348) and poly-A tailed (Poly(A) Tailing kit, ThermoFisher Scientific; Cat. No. AM1350) mRNA, in pre-warmed drops of M2 media overlaid with mineral oil, on the surface of concaved microscope slides. The non-targeting control siRNA (NTC) used was from Qiagen GeneGlobe (Qiagen, Hilden, Germany; Cat. No. SI03650318) and the *Ddx21* siRNA used was from Thermo Fisher Scientific Silencer Select (Thermo Fisher Scientific; Cat. No. 4390771 & assay ID: s80158).

### Immunofluorescence staining, confocal microscopy and image acquisition

To remove the *zona pellucida*, preimplantation embryos at the appropriate developmental stage were quickly washed and pipetted in pre-warmed drops of acidic Tyrode’s Solution (Sigma-Aldrich; Cat. No. T1788) until *zona pellucidae* were visually undetectable and then immediately washed through pre-warmed drops of M2 media. Embryos were fixed, in the dark, with 4% paraformaldehyde (Santa Cruz Biotechnology, Inc., Dallas, Texas, USA; Cat. No. sc-281692) for 20 minutes at room temperature. Permeabilisation was performed by transferring embryos to a 0.5% solution of Triton X-100 (Sigma-Aldrich; Cat. No. T8787), in phosphate buffered saline (PBS), for 20 minutes at room temperature. All washes post-fixation, permeabilisation and antibody staining were performed in PBS with 0.05% TWEEN 20 (Sigma-Aldrich; Cat. No. P9416) (PBST) by transferring embryos between two drops or wells (of 96-well micro-titre plates) of PBST, for 20 minutes at room temperature. Blocking and antibody staining was performed in 3% bovine serum albumin (BSA; Sigma-Aldrich; Cat. No. A7906) in PBST. Blocking incubations of 30 minutes at 4°C were performed before both primary and secondary antibody incubation; primary antibody incubation (in blocking buffer) was performed overnight (∼16 hours) at 4°C and secondary antibody incubation carried out in the dark at room temperature for 70 minutes. Stained embryos were mounted in DAPI containing mounting medium (VECTASHIELD; Vector Laboratories, Inc., Burlingame, California, USA; Cat. No. H-1200), placed under cover slips on glass-bottomed 35mm culture plates and incubated at 4°C for 30 minutes in the dark, prior to confocal imaging. Details of the primary and secondary antibody combinations used can be found in supplementary table S2. Complete embryo confocal z-series images were acquired using a FV10i Confocal Laser Scanning Microscope and FV10i-SW image acquisition software (Olympus, Tokyo, Japan). Images were analysed using FV10-ASW 4.2 Viewer (Olympus) and Imaris X64 Microscopy Image Analysis Software (version 6.2.1; Bitplane AG -Oxford Instruments plc, Abingdon, United Kingdom). Cells were counted both manually and semi-automatically using Imaris X64 encoded functions.

### Cell number quantification, statistics and graphical representation

Total embryo cell number counts (plus outer and inner cell populations) from confocal acquired z-series micrographs (based on DAPI nuclei staining) were further subcategorised as inner or outer based on absence or presence of CDX2 expression respectively (Fig. 3). EPI or PrE cells were quantified based on detectable and exclusive expression of NANOG and GATA4 protein expression, respectively (Fig. 4). Cells not located within blastocyst ICMs that also did not stain for either GATA4 and/or NANOG, were designated as outer/TE cells. Initial recording and data accumulation was carried out using Microsoft Excel and further statistical analysis and graphical representations were performed with GraphPad Prism 8. Based on the normality and lognormality comparisons, appropriate statistical tests were used for the compared datasets (summarised in supplementary table S3). Unless otherwise stated within individual graphs as a specific p-value (if statistically insignificant), the stated significance intervals were as follows: p-value < 0.0001 (****), 0.0001 to 0.001 (***), 0.001 to 0.01 (**) and 0.01 to 0.05 (*). All graphs represent dot plots of the total sample size, with associated means and standard deviation error bars provided.

### Fluorescence intensity quantification and statistical analysis

Levels of DDX21 (Fig. 2e, 3c-e and supplementary Fig. S2a and S3a-c) and NUCLEOSTEMIN (GNL3) protein (Fig. 2f and supplementary Fig. S2b) were quantified, using Fiji (ImageJ)^47^, as fluorescence intensity per cell nucleus and differentiated into inner and outer cells based on absence or presence of CDX2 immuno-fluorescence, respectively. The measurements settings for all of the above were as follows: Analyze>Set Measurements; and the following options were chosen: Area, Mean grey value and Integrated density. Using the “Polygon selections” tool, individual cell nuclei were demarcated and the aforementioned measurements recorded. The selected area was then moved so as to encompass an area excluding that of the embryo or cell nucleus, respectively, and background measurements were recorded. This process was continued for all the embryos analysed, under both control and p38-MAPKi conditions and the results were transferred to a spreadsheet for further calculations. The Corrected Total Cell Fluorescence (CTCF), in arbitrary units, for each embryo was measured as such: CTCF = Integrated Density – (Area of selected cell X Mean fluorescence of background readings)^48,49^ – supplementary tables S4 & 5. The calculated CTCF are plotted as scatters, with mean and standard deviations marked. The CTCF differences were statistically tested using appropriate statistical tests (supplementary table S3) based on normality and lognormality comparisons and the results, unless otherwise stated within individual graphs as a specific p-value (if statistically insignificant), are stated as following significance intervals: p-value < 0.0001 (****), 0.0001 to 0.001 (***), 0.001 to 0.01 (**) and 0.01 to 0.05 (*).

### Blastocyst size and volume calculations

Equatorial outer circumference and total volume measurements for the blastocyst as a whole were carried out by measuring the embryo outer circumference of the centrally located widest Z-stack using Fiji (ImageJ)^47^. The measurements were set as follows: Analyze>Set Measurements; and the “Perimeter” option was selected; using the “Polygon selections” tool, the outer circumference was traced and measured. The radius of the measured circumference was deduced and that value was used to calculate an approximate volume for all the embryos analysed under both control (NTC) and *Ddx21* siRNA injected conditions. The calculated volume in picoliters (pL) (tabulated in supplementary table S6 is plotted as a scatter, with mean and standard deviations marked. The differences were statistically tested using Mann-Whitney test and the results, unless otherwise stated within individual graphs as a specific p-value (if statistically insignificant), are stated at the following significance intervals: p-value < 0.0001 (****), 0.0001 to 0.001 (***), 0.001 to 0.01 (**) and 0.01 to 0.05 (*).

### Quantitative real-time PCR (qRT-PCR)

20 embryos per condition (control (NTC) and *Ddx21* siRNA injected) from global microinjection experiments, were collected at E4.5 and immediately processed for RNA extraction and isolation using the ARCTURUS PicoPure RNA Isolation Kit (Thermo Fisher Scientific; Cat. No. KIT0204), following the manufacturer’s protocol. The entire eluted volume of total RNA was immediately DNase treated with TURBO DNase ((Thermo Fisher Scientific; Cat. No. AM2238) according to the manufacturer provided protocol. The whole sample was then subject to cDNA synthesis using SuperScript III Reverse Transcriptase (Thermo Fisher Scientific; Cat. No. 18080044), as directed by the manufacturer and employing oligo d(T)16 (Thermo Fisher Scientific; Cat. No. N8080128), dNTP Mix (Thermo Fisher Scientific; Cat. No. R0192) and RNase Inhibitor (Thermo Fisher Scientific; Cat. No. N8080119). The synthesized cDNA was diluted as required with nuclease free water and 1µl was used in 10µl individual final SYBR-green based qRT-PCR reaction volumes (PCR Biosystems Ltd., London, United Kingdom; Cat. No. PB20.11) with oligonucleotide primers at a final concentration of 300nM each. A Bio-Rad CFX96 Touch Real-Time PCR Detection System apparatus, employing standard settings, was used for data accumulation and initial analysis was performed with the accompanying Bio-Rad CFX Manager software. Triplicate measurements per gene were assayed from one experiment that were each technically replicated. The sequence of oligonucleotide primers (5’ to 3’) used were: i) *Tbp* -GAAGAACAATCCAGACTAGCAGCA (sense) and CCTTATAGGGAACTTCACATCACAG (anti-sense) and ii) -*Ddx21*: TTCCTTCTGCAACGGAAATAA (sense) and GAGGCACAGAATCCAAGAGC (anti-sense). The average transcript levels of the *Ddx21* gene were derived after internal normalisation against *Tbp* mRNA levels. Data was acquired and initially analysed with CFX Manager, then processed in Microsoft Excel (biological and technical replicate averaging) and GraphPad Prism 9 (graphical output).

## Supplementary figure legends

**1. Figure S1: Gene expression profile of *Ddx21*** Gene expression profile of *Ddx21* mRNA in oocyte, preimplantation embryos and early embryonic cell lineages. Data collected from (a) *Zhang, B*., *Zheng, H*., *Huang, B. et al. Allelic reprogramming of the histone modification H3K4me3 in early mammalian development. Nature 537, 553–557 (2016)* and (b) *Wang, C*., *Liu, X*., *Gao, Y. et al. Reprogramming of H3K9me3-dependent heterochromatin during mammalian embryo development. Nat Cell Biol 20, 620–631 (2018)*.

**2. Figure S2: Supplementary to Figure 2. Effect of p38-MAPK inhibition on DDX21 and NUCLEOSTEMIN (GNL3) protein expression**. Comparing per nucleus corrected fluorescence (CTCF) of (a) DDX21 and (b) NUCLEOSTEMIN (GNL3) between inner and outer cells (based on CDX2 expression) at E4.5 after 24 hours (from E3.5) of treatment in both control (DMSO) and p38-MAPKi (SB220025) conditions.

**3. Figure S3: Supplementary to Figure 3. Clonal knockdown of *Ddx21* and effect on late-blastocyst morphology and cell numbers**. Scatter plot quantification of per cell CTCF of DDX21 expression in control (NTC siRNA injected n=15 embryos; non-injected cells outer n=151 & inner n=108, injected cells outer n=153 & inner n=112) and *Ddx21* downregulated (*Ddx21* siRNA injected n=26; non-injected cells outer n=225 & inner n=125, injected cells outer n=157 & inner n=59) embryos, a) Combined for cells from embryos injected with NTC and *Ddx21* siRNA b) Only outer cells, comparing injected and non-injected cell clones between NTC and *Ddx21* siRNA. c) Only inner cells, comparing injected and non-injected cell clones s between NTC and *Ddx21* siRNA. d) Scatter plot quantification of cell numbers comparing NTC siRNA and *Ddx21* siRNA microinjected embryos, categorised between microinjected, non-microinjected clones and combined cell number and further divided between (i) total, (ii) outer and (iii) inner cell populations.

**4. Figure S4: Supplementary to Figure 4. Effect of *Ddx21* knockdown on Epiblast (EPI) and Primitive Endoderm (PrE) lineage specification at E4.5**. Scatter plot quantification of (i) total cell numbers (DAPI nuclear stain), (ii) outer cell numbers, (iii) inner cell numbers, (iv) ratio of EPI to ICM, (v) number of NANOG-GATA4 co-negative inner cells and (vi) ratio of NANOG-GATA4 co-negative inner cells to ICM of clonal NTC siRNA and *Ddx21* siRNA microinjected injected embryos, as observed in either the microinjected or non-injected clones and both clones combined.

## Supplementary table legends

**1. Table S1: Phosphoproteomic mass spectrometry result regarding detection of DDX21 in DMSO (control) vs. p38-MAPKi (SB220025) conditions**.

**2. Table S2: List of antibodies used**.

**3. Table S3: List of statistical tests used for respective data analysis**.

**4. Table S4: Collated individual nuclei CTCF quantifications for DDX21 and NUCLEOSTEMIN (GNL3) in DMSO (control) vs. p38-MAPKi (SB220025) conditions**. (Supplementary to Fig. 2e, f & S2a, b)

**5. Table S5: Collated individual nuclei CTCF quantifications for DDX21 in control (NTC siRNA injected) and Ddx21 downregulated embryos (Ddx21 *siRNA*)**. (Supplementary to Fig. 3c-e & S3a-c)

**6. Table S6: Collated individual embryo volume (in picoliter (pL))**. (Supplementary toFig. 3j)

**7. Table S7: Collated individual embryo cell number quantifications in control (NTC siRNA injected) and Ddx21 downregulated embryos (*Ddx21 siRNA)***. Cells are divided into inner and outer based on absence or presence of CDX2 expression respectively and non-injected or injected based on absence or presence of RFP respectively (Supplementary to Fig. 3f-i & S3d).

**8. Table S8: Collated individual embryo cell number quantifications in control (NTC siRNA injected) and Ddx21 downregulated embryos (*Ddx21 siRNA)***. Cells are divided based on physical location (inner and outer) and on NANOG and GATA4 expression. Non-injected or injected demarcations are based on absence or presence of RFP respectively (Supplementary to Fig. 4 & S4).

## Acknowledgements

We acknowledge the Institute of Parasitology (Biology Centre of the Czech Academy of Sciences, in České Budějovice) for housing mice, Marta Gajewska (Institute of Oncology, Warsaw, Poland) and Anna Piliszek (Institute of Genetics and Animal Breeding, Polish Academy of Sciences, Jastrzębiec, Poland) for founder CBA/W mice, Alena Krejčí (Faculty of Science, University of South Bohemia, Czech Republic) for pooling resources and other members of our laboratory for valuable inputs and discussions. We also acknowledge the core facility Masaryk University, Brno, Czech Republic -Faculty of Informatics, supported by the MEYS CR (LM2018129 Czech-BioImaging) for assistance with image analysis. CIISB, Instruct-CZ Centre of Instruct-ERIC EU consortium, funded by MEYS CR infrastructure project LM2018127, is gratefully acknowledged for the financial support of the measurements at the CEITEC Proteomics Core Facility.

