## Supplementary figures for "DDX21 is a p38-MAPK sensitive nucleolar protein necessary for mouse preimplantation embryo development and cell-fate specification"

Fig S1

a Gene expression profile of *Ddx21* from *Zhang et al. Nature 537, 553–557 (2016)*

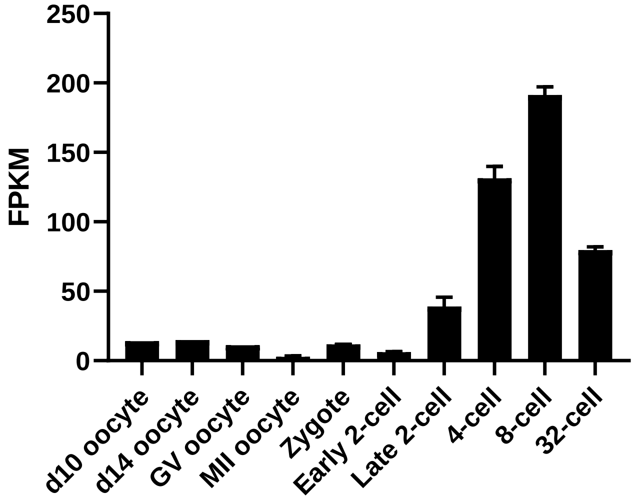


b Gene expression profile of *Ddx21* from *Wang et al. Nat Cell Biol 20, 620–631 (2018)*

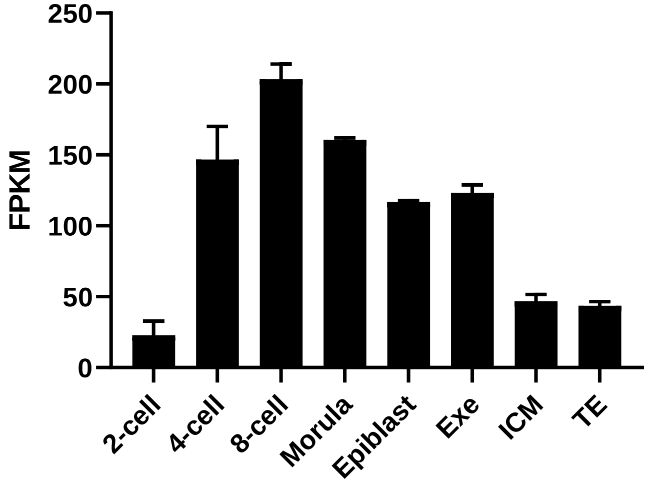


Gene expression profile of *Ddx21* in oocyte, preimplantation embryos and early embryonic cellular lineages. Data collected from *Zhang, B., Zheng, H., Huang, B. et al. Allelic reprogramming of the histone modification H3K4me3 in early mammalian development. Nature 537, 553–557 (2016)* and *Wang, C., Liu, X., Gao, Y. et al. Reprogramming of H3K9me3-dependent heterochromatin during mammalian embryo development. Nat Cell Biol 20, 620–631 (2018)*.

### Fig S2

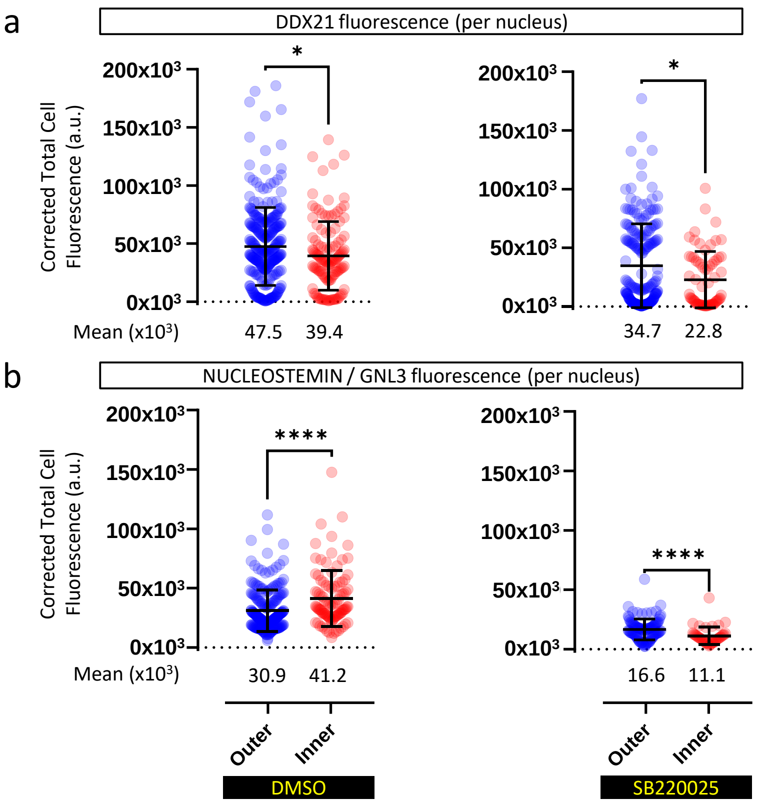


Supplementary to Figure 2. Effect of p38-MAPK inhibition on DDX21 and NUCLEOSTEMIN (GNL3) protein expression.

Comparing per nucleus corrected fluorescence (CTCF) of (a) DDX21 and (b) NUCLEOSTEMIN (GNL3) between inner and outer cells (based on CDX2 expression) at E4.5 after 24 hours (from E3.5) of treatment in both control (DMSO) and p38-MAPKi (SB220025) conditions.

### Fig S3

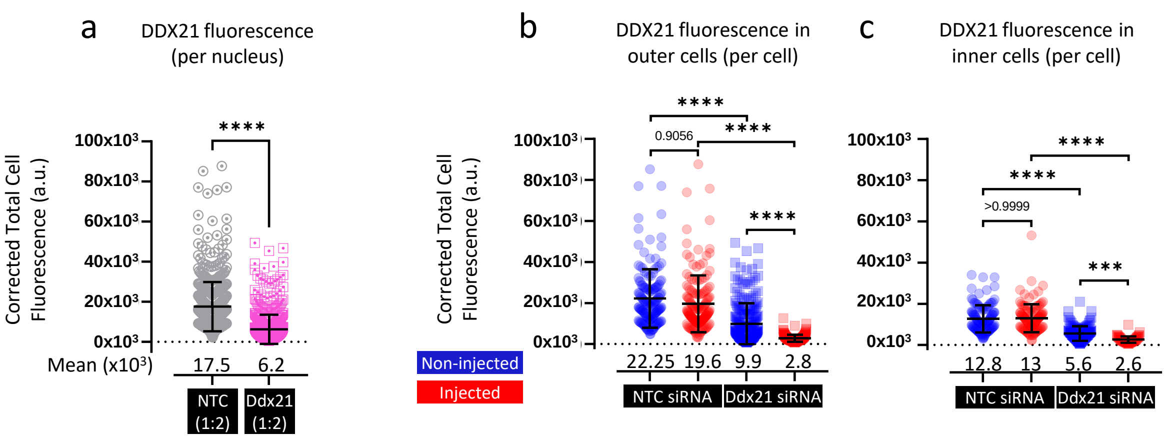


Supplementary to Fig. 3c-e.

Scatter plot quantification of per cell CTCF of DDX21 expression in control (NTC siRNA injected n=15 embryos; non-injected cells outer n=151 & inner n=108, injected cells outer n=153 & inner n=112) and Ddx21 downregulated (Ddx21 siRNA injected n=26; non-injected cells outer n=225 & inner n=125, injected cells outer n=157 & inner n=59) embryos,

a) Combined for cells from embryos injected with NTC and Ddx21 siRNA

b) Only outer cells, comparing injected and non-injected cell clones between NTC and Ddx21 siRNA.

c) Only inner cells, comparing injected and non-injected cell clones s between NTC and Ddx21 siRNA.

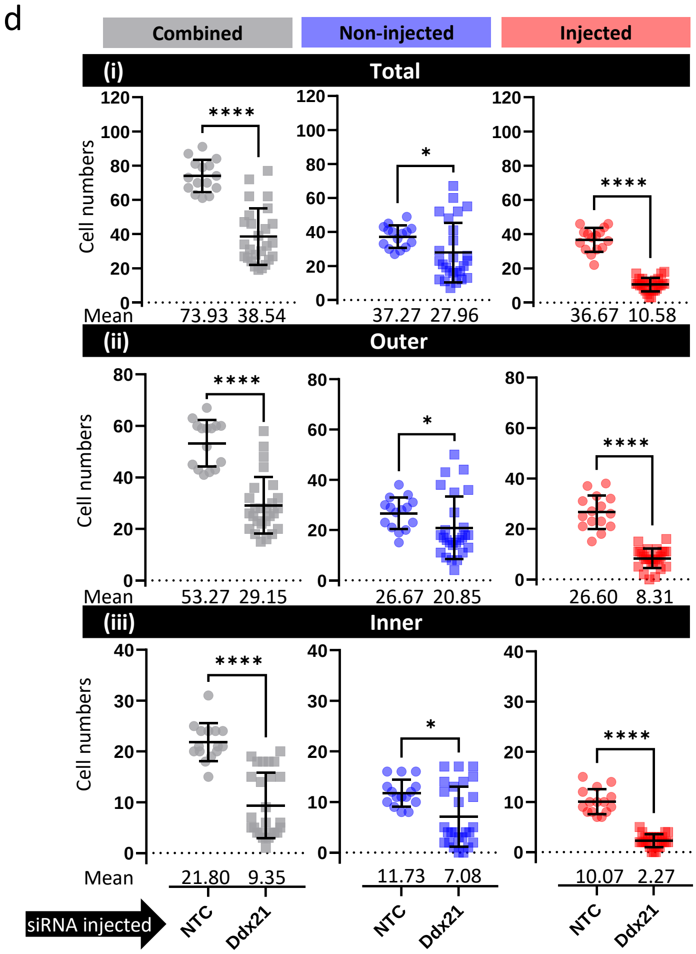


Supplementary to Fig. 3h, i.

d) Scatter plot quantification of cell numbers comparing NTC siRNA and Ddx21 siRNA microinjected embryos, categorised between microinjected, non-microinjected clones and combined cell number and further divided between (i) total, (ii) outer and (iii) inner cell populations.

### Fig S4

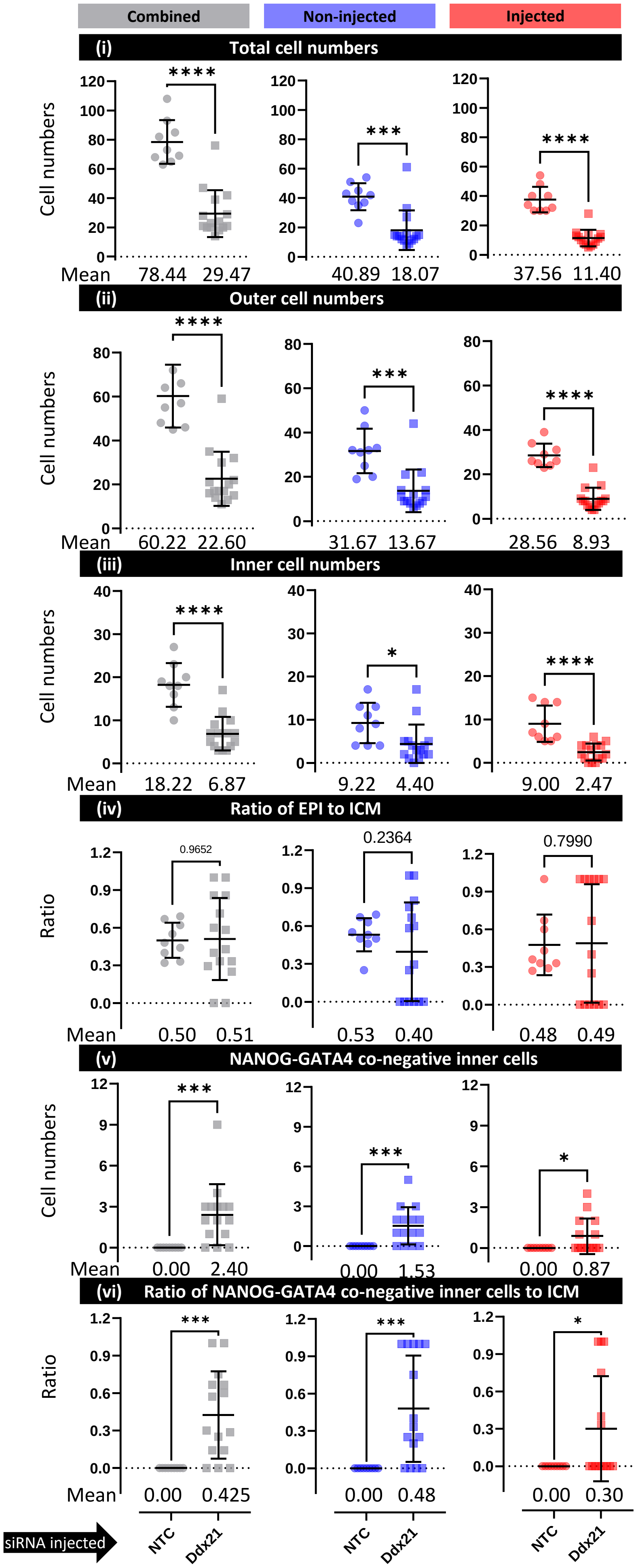


Supplementary to Fig. 4

Scatter plot quantification of (i) total cell numbers (DAPI nuclear stain), (ii) outer cell numbers, (iii) inner cell numbers, (iv) ratio of EPI to ICM, (v) number of NANOG-GATA4 co-negative inner cells and (vi) ratio of NANOG-GATA4 co-negative inner cells to ICM of clonal NTC siRNA and Ddx21 siRNA microinjected injected embryos, as observed in either the microinjected or non-injected clones and both clones combined.
